# Digit evolution in gymnophthalmid lizards

**DOI:** 10.1101/013953

**Authors:** Juliana G. Roscito, Pedro M.S Nunes, Miguel T. Rodrigues

**Affiliations:** JGR, MTR: Departamento de Zoologia, Instituto de Biociências, Universidade de São Paulo-SP; PMSN: Departamento de Zoologia, Centro de Ciências Biológicas, Universidade Federal de Pernambuco

**Keywords:** limb reduction, reptiles, morphological evolution, limb development

## Abstract

**Background:** The tetrapod limb is a highly diverse structure, and reduction of the limbs accounts for much of the phenotypes observed within species. Squamate reptiles represent one of the many lineages in which the limbs have been greatly modified from the pentadactyl generalized pattern; within the group, limb-reduced morphologies can vary from minor reductions in size of elements to complete limblessness, with several intermediate forms in between. Even though limb reduction is widespread, it is not clear what are the evolutionary and developmental mechanisms involved in the formation of reduced limb morphologies.

**Methods:** In this study, we present an overview of limb morphology within the microteiid lizard group Gymnophthalmidae, focusing on digit number.

**Results:** We show that there are two major groups of limb-reduced gymnophthalmids. The first group is formed by lizard-like - and frequently pentadactyl - species, in which minor reductions (such as the loss of 1-2 phalanges mainly in digits I and V) are the rule; these morphologies generally correspond to those seen in other squamates. The second group is formed by species showing more drastic losses, which can include the absence of an externally distinct limb in adults. We also show the expression patterns of Sonic Hedgehog (Shh) in the greatly reduced fore and hindlimb of a serpentiform gymnophthalmid.

**Conclusions:** Our discussion focus on identifying shared patterns of limb reduction among tetrapods, and explaining these patterns and the morphological variation within the gymnophthalmids based on the current knowledge of the molecular signaling pathways that coordinate limb development.

## Introduction

The tetrapod limb has been the subject of extensive investigation in evolutionary and developmental biology for over a century. Since the 19^th^ century, the evolutionary origin and diversification of the limbs have been greatly debated, and several hypothesis and mechanisms have been proposed to explain how such diversity arose during evolution.

Even though the identification of evolutionary patterns of limb evolution are crucial, the fundamental question of how does such diversity arises is better answered by studying the mechanisms involved in the formation of the limbs. This is not a trivial investigation, because it involves getting access to embryonic material, and being able to experimentally manipulate such embryos, only possible with a more in depth knowledge of the genetics of the organism.

The bulk of knowledge on limb development in tetrapods relies, up to the moment, on the two most extensively studied organisms, the lab mouse and chicken. The signaling networks controlling limb development seem to be fairly conserved among those species, although important differences exist. Great reviews have been published summarizing the current knowledge of how limb development is organized (Niswander, 2003; Zeller *et al*., 2009; Rabinowitz and Vokes, 2012); for a more detailed picture of such mechanisms, the reader should refer to that literature and references within. However, variations of those mechanisms among natural populations should be expected, and have, in fact, been demonstrated for some species (Thewissen *et al*., 2006; Cooper *et al*., 2007; Hockman *et al*. 2008; Doroba and Sears, 2010; Sears *et al*., 2011; Cooper *et al*. 2014; Lopez-Rios *et al*., 2014).

A group that is underrepresented in developmental studies is the Squamata. Understanding the evolution and diversification of the reptile limb has been the focus of studies in several research fields, including evolutionary biology, paleontology, ecology, anatomy, physiology, functional morphology and, to a lesser degree, developmental biology (Gans, 1975; Lande, 1978; Withers, 1981; Greer, 1991; Caldwell, 2002; Shapiro, 2002; Shapiro *et al*., 2003; Whiting *et al*., 2003; Crumly and Sanchez-Villagra, 2004; Kearney and Stuart, 2004; Kohlsdorf and Wagner, 2006; Wiens *et al*., 2006, Brandley *et al*., 2008; Kohlsdorf *et al*., 2008; Russel and Bauer, 2008; Skinner *et al*., 2008; Bergmann and Irschick, 2009; Jerez and Tarazona, 2009; Young *et al*., 2009; Leal *et al*., 2010; Hugi *et al*., 2012; etc).

A recurrent phenotype among squamate reptiles is limb reduction. From minor losses of phalanges, to the complete loss of the limb, there is a wide spectrum of intermediate morphologies that can occur even among species that belong to the same genus (Choquenot and Greer, 1989; Skinner *et al*., 2008). Changes like those are extremely frequent, having occurred multiple times independently in almost all major squamate groups (Greer, 1991; Wiens *et al*. 2006; Skinner *et al*., 2008). The scincids and anguids top the rank of limb-reduced lineages, but reduction is also seen in the pygopodids, gekkonids, cordylids, gerrhosaurids, dibamids, gymnophthalmids, amphisbaenians, and, obviously, in the snakes, which have attained the greatest degree of limb reduction. Previous studies have successfully identified patterns of limb reduction among squamates and provided evolutionary, functional, anatomical, and biogeographical/environmental hypothesis for the evolution of reduced forms (Greer, 1991; Benesch and Withers, 2002; Wiens *et al*., 2006; Brandley *et al*., 2008; Camacho *et al*., 2014).

Even though an incredible diversity of limbs is observed among squamate reptiles (lizards, snakes and amphisbaenians), little is known about the mechanisms involved in limb development within the group. Most of the studies concern morphological analyses of limb development, which are important for laying the anatomical foundations for further functional investigations (for example, Howes and Swinnerton, 1901; Mathur and Goel, 1976; Rieppel, 1994; Shapiro, 2002; Fabrezi *et al*., 2007; Leal *et al*., 2010; Roscito and Rodrigues, 2012a,b), but still few studies (Raynaud, 1990; Raynaud *et al*., 1998; Cohn and Tickle, 1999; Shapiro *et al*., 2003; Young *et al*., 2009) attempted to undercover the molecular mechanisms behind limb development in natural populations of squamate species.

Among the Gymnophthalmidae, a South American group of microteiid lizards, limb-reduced species are widespread, occurring in desert-like environments, highlands, and also forested areas. Despite the diversity of forms, we still lack a clear picture of the diversity of limb morphologies within the group, how are such morphologies distributed across the lineages, and how could they have evolved. Thus, in this paper we present an extensive survey of the limbs of several gymnophthalmid species with the aim of identifying patterns of limb reduction within the group. We also present a preliminary investigation of the molecular signaling involved in the formation of a reduced limb, and analyse the different types of reduction based on the current knowledge of the mechanisms controlling limb development in vertebrates.

## METHODS

### Material examined

Our sample is comprised of 34 Gymnophthalmid genera and 70 species, out of a total of 46 genera and 244 species currently recognized for the group (reptile-database.org; august/2014). We included in our sampling only those species for which we could get information of the phalangeal formula, either by observing cleared and stained material, or by information from the literature.

Cleared and double stained specimens are from our personal collection and from the collection of Museu de Zoologia da Universidade de São Paulo; the material examined and respective collection numbers is listed in the **supplementary information**.

The notation used for representing the phalangeal formula is the following:

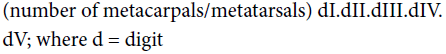

### Phylogenies of the Gymnophthalmidae

Up to the present, there are three phylogenies for the Gymnophthalmidae (Pellegrino *et al*., 2001; Castoe *et al*., 2004; Pyron *et al*., 2013). The same species are consistently grouped together in all three phylogenies (Figure 1). However, each author assigns each group to differently inclusive taxonomic rankings, which generates incongruences over the status of particular gymnophthalmid groups (see discussion in Rodrigues *et al*., 2007b, 2009). In this paper we do not wish to solve those incongruences, therefore our naming of the major groups and subgroups recognized within the Gymnophthalmid reflects our personal ideas regarding the evolution of these species:

**Figure 1.**
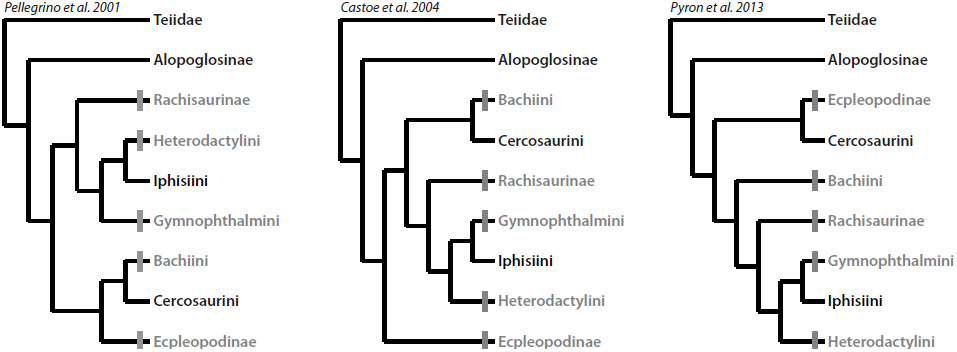
Relationships within the Gymnopthalmidae, adapted from the phylogenetic hypothesis of Pellegrino *et al*. (2001), Castoe *et al*. (2004), and Pyron *et al*. (2013). Lineages with limb-reduced species are marked in grey.

**Alopoglossinae** (Pellegrino *et al*., 2001; Castoe *et al*., 2004): *Alopoglossus, Ptychoglossus*

**Rachisaurinae** (Pellegrino *et al*., 2001; Castoe *et al*., 2004): *Rachisaurus*

### Cercosaurinae

**Bachiini** (Castoe *et al*., 2004): *Bachia*

**Cercosaurini** (Castoe *et al*., 2004): *Cercosaura, Echinosaura*, *Neusticurus*, *Placosoma*, *Pholidobolus*, *Potamites, Proctoporus*.

**Ecpleopodinae** (Castoe *et al*., 2004; Pyron *et al*., 2013): *Anotosaura, Arthrosaura, Colobosauroides, Dryadosaura, Leposoma*.

### Gymnophthalminae

**Heterodactylini** (Rodrigues *et al*., 2009): *Caparaonia, Colobodactylus, Heterodactylus*

**Iphisiini** (Rodrigues *et al*., 2009): *Acratosaura, Al-exandresaurus, Colobosaura, Iphisa, Stenolepis*

**Gymnophthalmini** (Pellegrino *et al*., 2001): *Tretioscincus, Gymnophthalmus, Micrablepharus, Procellosaurinus, Psilophthalmus, Vanzosaura, Nothobachia, Scriptosaura, Calyptommatus*

### Embryonic material and whole mount in-situ

We obtained a small developmental series of five *Calyptommatus sinebrachiatus* embryos during a field trip to Bahia State, Brazil, in 2005, that were fixed in RNAlater after 6, 10, 12, 14, and 16 days after laying. The whole-mount in-situ hybridization for detection of the gene *Sonic hedgehog* followed a modified protocol used for chicken embryos, starting with two 5-minute washes in 100% methanol, with a 1-hour in between incubation in 6% hydrogen peroxide solution (in 100% methanol). Following rehydration of the embryos in PBT, the embryos were subjected to a proteinase K digestion (20 minute reaction at room temperature) followed by re-fixation in 4% PFA for 20 minutes. Embryos then went through to a series of ‘pre-probe’ incubations in hybridization buffer and finally incubated overnight at 65oC with a chicken-specific probe for Shh (1:10 dilution). After a series of washes with MABT, and a blocking step, embryos were incubated overnight with anti-DIG (1:2000), and then the signal was revealed with NBT/BCIP.

## RESULTS

### Digit arrangement in gymnophthalmid lizards

The phalangeal formula of both fore and hindlimb, as well as specific comments on the morphology of metacarpal/ metatarsal and phalangeal elements, are presented in Table 1.

**Table 1.** Summary of the phalangeal formulas for Gymnophthalmidae species. Entries in grey correspond to those species not directly analyzed. dc/dt, distal carpal/distal tarsal; mc/mt, metacarpal/metatarsal.

| Species | Hand phalan-geal formula | Foot phalan-geal formula | Observations | References |
| --- | --- | --- | --- | --- |
| *Acratosaura mentalis* | (5) 2:3:4:5:3 | (5) 2:3:4:5:4 | Both fore and hindlimb follow the pentadactyl condition. |  |
| *Alexandresaurus camacan* | (5) 2:3:4:5:3 | (5) 2:3:4:5:4 | Both fore and hindlimb follow the pentadactyl condition, but the last phalanx (ungual phalax) from digit I in the hand is shorter and ends in a relatively blunt end in comparison to the other four ungual phalanges, which are longer and sharper. Rodrigues et al. (2007) mention that digit I from forelimb bears no claw. | *Rodrigues et al. (2007)* |
| *Alopoglossus angulatus* | (5) 2:3:4:5:3 | (5) 2:3:4:5:4 | Both fore and hindlimb follow the pentadactyl condition. |  |
| *Alopoglossus atriventris* | (5) 2:3:4:5:3 | (5) 2:3:4:5:4 | Both fore and hindlimb follow the pentadactyl condition. Phalangeal formula following Grizante (2009). | *Grizante (2009)* |
| *Anotosaura collaris* | (5) 2:3:4:4:3 | (5) 2:3:4:5:3 | Osteology of the limbs described in Rodrigues et al. (2013); Kizirian and McDiarmid (1998) agree on the phalan-geal formula of the hindlimb.<br>The 5th phalanx from digit IV of the forelimb, and the 4th phalanx of digit V of the hindlimb, are absent. | *Kizirian and McDiarmid (1998);*<br>*Rodrigues et al. (2013)* |
| *Anotosaura vanzolinia* | (5) 2:3:4:4:3 | (5) 2:3:4:5:4 | Forelimb is short and stout, and the 5th phalanx from digit IV is absent. The same condition is reported by Kizirian and McDiarmid (1998). | *Kizirian and McDiarmid (1998)* |
| *Arthrosaura hoogmoedi* | (5) 2:3:4:5:3 | (5) 2:3:4:5:4 | Both fore and hindlimb follow the pentadactyl condition. No osteological data presented in Kok (2008). | *Kok (2008)* |
| *Arthrosaura kockii* | (5) 2:3:4:5:3 | (5) 2:3:4:5:4 | Both fore and hindlimb follow the pentadactyl condition. Phalangeal formula following Grizante (2009). | *Grizante (2009)* |
| *Arthrosaura reticulata* | (5) 2:3:4:5:3 | (5) 2:3:4:5:4 | Both fore and hindlimb follow the pentadactyl condition. |  |
| *Bachia barbouri* | (2) 1?:1? | (?) 0:0:0:0:0 | Kizirian and McDiarmid (1998) did not determined the identity of the digits in the forelimb.<br>Hindlimbs are absent or tubercular (Dixon, 1973).<br>Kohlsdorf and Wagner (2006) reported the phalangeal formulas of 0:0:2:2:0 for the forelimb and 0:0:0:0:0 for the hindlimb. | *Kizirian and McDiarmid (1998);*<br>*Dixon (1973); Kohlsdorf and*<br>*Wagner (2006)* |
| *Bachia bicolor* | (4) 0:2:2:2:2 | (0) 0:0:0:0:0 | Phalangeal formula following Kizirian and McDiarmid (1998) and Jerez and Tarazona (2009). Thomas (1965) states the presence of 1 or 2 digits in the hindlimb. | *Kizirian and McDiarmid (1998);*<br>*Jerez and Tarazona (2009);*<br>*Thomas (1965)* |
| *Bachia blairi* | (5) 0:2:2:2:2 | (5) 2:2:2:0:0 | Phalangeal formula following Kizirian and McDiarmid (1998). | *Kizirian and McDiarmid (1998)* |
| *Bachia bresslaui* | (2) 0:0:1?:2?:0 | (?) 0:0:0:0:0 | Phalangeal formula following Kizirian and McDiarmid (1998). One cleared and double stained specimen was available for analysis, but the forelimbs and right hindlimb were damaged; in the left hindlimb we could not identify the elements distal to tibia/fibula. | *Kizirian and McDiarmid (1998)* |
| *Bachia dorbignyi* | (4) 0:2:2:2:0 | (?) 0:0:0:0:0 | Phalangeal formula following Kizirian and McDiarmid (1998). | *Kizirian and McDiarmid (1998)* |
| *Bachia flavescens* | (5) 0:1:1:1:0 | (3) 0:0:0:0:0 | Due to poor condition of the specimen available for analysis, we cannot affirm that both fore and hindlimb were not damaged.<br>Forelimb with five metacarpals and one phalanx in each of the digits II, III, and IV.<br>The hindlimb was either less preserved, or not properly stained: the left hindlimb had only the femur, while in the right limb the zeugopodial elements (tibia/fibula) were clearly visible. The autopod was faintly stained, so we believe that the digits likely correspond to digits III, IV, and V, because of their position distal to the fibula. Our observations agree with Thomas (1965), which also reported three digits in the forelimb and no digits in the hindlimb of *Bachia parkeri* (which is now synonymized to *B. flavescens*, according to *reptile-database.com*). Kizirian and McDiarmid (1998), on the other hand, reported the phalangeal formula of (4) 0:2:2:2:0 and (4) 1:0:0:0:0 for fore and hindlimb, respectively, for *Bachia flavescens parkeri*. | *Kizirian and McDiarmid (1998); Thomas (1965)* |
| *Bachia heteropa* | (5) 1:2:3:3:2 | (5) 2:2:3:3:0 | Phalangeal formula following Kizirian and McDiarmid, 1998. | *Kizirian and McDiarmid (1998)* |
| *Bachia huallagana* | (?)/3 digits | 0:0:0:0:0 | Number of digits in the forelimb following Dixon (1973), but the description is based only on external morphology. The author mentions that hindlimb is “tubercular”. Kizirian and McDiarmid (1998) only observed the hindlimb, thus, the phalangeal formula is derived from their report. | *Kizirian and McDiarmid (1998); Dixon (1973)* |
| *Bachia intermedia* | (4) 0:2:2:2:0 | (?) 0:0:0:0:0 | Phalangeal formula following Kizirian and McDiarmid, 1998. Thomas (1965) agrees with Kizirian and McDiarmid (1998) regarding the presence of 3 digits in the forelimb, but reports the presence of 2 digits in the hindlimb; his analysis rely only on external morphology. Dixon (1973) mentions that hindlimb is styliform “with one or two apical scales resembling toes”. Presch (1975) reports that tibia and fibula are fused in this species. | *Kizirian and McDiarmid (1998); Thomas (1965); Dixon (1973); Presch (1975)* |
| *Bachia monodactylus monodactylus* | (?) 0:1:1:1:1 | (?)/1 digit? | Phalangeal formula following Kohlsdorf and Wagner (2006) | *Kohlsdorf and Wagner (2006)* |
| *Bachia pallidiceps* | (5) 0:2:2:2:2 | (5) 2:2:2:0:0 | Phalangeal formula following Kizirian and McDiarmid (1998). | *Kizirian and McDiarmid (1998)* |
| *Bachia panoplia* | (4/5) 0:2:2:2:2 | (4) 2:2:2:2:0 | The hand plate has four metacarpals (or five: a small nodular element posterior to dc II and metacarpal II can be either identified as mcl or dc I; likely, Kizirian and McDiarmid (1998) considered the nodular element as mcl), and two phalanges in each of the digits II-V. Digits I-IV of the hindlimb are composed of a metatarsal and two phalanges each. |  |
| *Bachia peruana* | (?) 3 digits | (?) 0:0:0:0:0 | Number of digits and phalangeal formula following Kizirian and McDiarmid (1998). The authors observed only the hindlimb. | *Kizirian and McDiarmid (1998)* |
| *Bachia pyburni* | (5) 0:3:4:4:3 | (5) 2:3:4:4:0 | Phalangeal formula following Kizirian and McDiarmid (1998). Rodrigues et al. (2008) agree regarding the number of digits present in both fore and hindlimb, although no osteological data was presented by the later. | *Kizirian and McDiarmid (1998); Rodrigues et al. (2008)* |
| *Bachia scolecoides* | (5) 0:2:2:2:2 | (4) 2:2:2:2:0 | Phalangeal formula following Kizirian and McDiarmid (1998) Rodrigues et al. (2008) agree regarding the number of digits present in both fore and hindlimb, although no osteological data was presented by the later. | *Kizirian and McDiarmid (1998); Rodrigues et al. (2008)* |
| *Bachia trisanale* | (3) 0:0:1:1:0 | (?) 0:0:0:0:0 | Phalangeal formula following Kizirian and McDiarmid (1998). | *Kizirian and McDiarmid (1998)* |
| *Calyptommatus leiolepis*<br>*Calyptommatus nicterus*<br>*Calyptommatus sinebrachiatus* | humerus | dIV | Osteology of the limbs described in Roscito and Rodrigues (2013).<br>Forelimb is absent externally, formed by a rudimentary humerus which does not protrude from the body wall. The single digit of the hindlimb is formed by one metatarsal and two phalanges (the last one being the ungual phalanx and bearing a nail). | *Roscito and Rodrigues (2013)* |
| *Caparaonia itaquara* | (5) 2:3:4:5:3 | (5) 2:3:4:5:4 | Both fore and hindlimb follow the pentadactyl condition, but the last phalanx (ungual phalax) from digit I of the forelimb is shorter and ends in a relatively blunt end in comparison to the other four ungual phalanges, which are longer and sharper. Rodrigues et al. (2009) note that digit I from forelimb bears no claw. | *Rodrigues et al. (2009)* |
| *Cercosaura eigenmanni* | (5) 2:3:4:5:3 | (5) 2:3:4:5:3 | Both fore and hindlimb follow the pentadactyl condition. |  |
| *Cercosaura ocellata* | (5) 2:3:4:5:3 | (5) 2:3:4:5:4 | Both fore and hindlimb follow the pentadactyl condition. |  |
| *Cercosaura schreibersii* | (5) 2:3:4:5:3 | (5) 2:3:4:5:4 | Both fore and hindlimb follow the pentadactyl condition. |  |
| *Colobodactylus dalcyanus* | (5) 1:3:4:5:3 | (5) 2:3:4:5:4 | Digit I from the forelimb is formed by a metacarpal and an extremely reduced phalanx, which does not look like an ungual phalanx. Contrary to our observations, Kizirian and McDiarmid (1998) and Grizante (2009) reported absence of phalanges in this same digit of the forelimb [(5) 0:3:4:5:3]. | *Kizirian and McDiarmid (1998); Grizante (2009)* |
| *Colobodactylus taunayi* | (5) 0/1:3:4:5:3 | (5) 2:3:4:5:4 | Digit I of the forelimb from one of the specimens examined (MTR 00746) shows a single phalanx following metacarpal I in both forelimbs (Fig. 3B). The other specimen analysed shows an almost imperceptible ossification distal to metacarpal I which can correspond to the single phalanx (Fig. 3C). Grizante (2009) reports no phalanges in digit I of the forelimb. | *Grizante (2009)* |
| *Colobosaura modesta* | (5) 2:3:4:5:3 | (5) 2:3:4:5:4 | Both phalanges from digit I of the forelimb are reduced, and the ungual phalanx doesn't end in a nail. Kizirian and McDiarmid (1998) and Grizante (2009) agree with our observations regarding phalangeal formulas. | *Kizirian and McDiarmid (1998); Grizante (2009)* |
| *Colobosauroides cearensis* | (5) 2:3:4:5:3 | (5) 2:3:4:5:4 | Both fore and hindlimb follow the pentadactyl condition. In specimen MZUSP 79595 we observed an asymmetry in phalange number between left and right forelimbs (4 or 5 phalanges in digit IV). |  |
| *Dryadosaura nordestina* | (5) 2:3:4:4:3 | (5) 2:3:4:5:4 | Forelimb digits are short and the last phalanx of digit IV is missing. Hindlimb follows the pentadactyl condition. |  |
| *Echinosaura horrida* | (5) 2:3:4:5:3 | (5) 2:3:4:5:4 | Both fore and hindlimb follow the pentadactyl condition. |  |
| *Gymnophthalmus underwoodi* | (5) 1:3:4:5:3 | (5) 2:3:4:5:4 | Digit I of the forelimb is short, showing a single reduced phalanx. Hindlimb follows the pentadactyl condition. |  |
| *Heterodactylus imbricatus* | (5) 1:3:4:5:3 | (5) 2:3:4:5:4 | There are minor reductions in phalange number in both fore and hindlimb. The single phalanx from digit I of the forelimb is reduced and does not resemble an ungual phalanx. The hindlimb follows the pentadactyl condition, although Kizirian and McDiarmid (1998) report only three phalanges in digit V [(5)2:3:4:5:3] whereas we observed 4; Grizante (2009) agrees with our observation on phalangeal formula. | Kizirian and McDiarmid (1998);<br>Grizante (2009) |
| *Heterodactylus lundii* | (5) 1/2:3:4:5:3 | (5) 2:3:4:5:4 | Forelimb has a reduced digit I. We observe an asymmetry between the left and right forelimbs, which show either one or two very small phalanges in this digit; the second phalanx seems vestigial. Hindlimb follows the pentadactyl condition.<br>Kizirian and McDiarmid (1988) report a single phalanx in digit I of the forelimb, and also an asymmetry between left and right hindlimbs, which show either 3 or none phalanges in digit V [(5) 2:3:4:5:3/0]. | Kizirian and McDiarmid (1998) |
| *Iphisa elegans* | (5) 2:3:4:5:3 | (5) 2:3:4:5:4 | Both fore and hindlimb follow the pentadactyl condition. However, the two phalanges of digit I of the forelimb are very reduced and the last one does not resemble an ungual phalanx. |  |
| *Leposoma guianensis* | (5) 2:3:4:5:3 | (5) 2:3:4:5:4 | Both fore and hindlimb follow the pentadactyl condition. Grizante (2009) agrees with our observation on the phalangeal formula. | Grizante (2009) |
| *Leposoma osvaldoi* | (5) 2:3:4:5:3 | (5) 2:3:4:5:4 | Both fore and hindlimb follow the pentadactyl condition. |  |
| *Leposoma percarinatum* | (5) 2:3:4:5:3 | (5) 2:3:4:5:4 | Both fore and hindlimb follow the pentadactyl condition. Grizante (2009) agrees with our observation on the phalangeal formula. | Grizante (2009) |
| *Micrablepharus atticolus*<br>*Micrablepharus maximiliani* | (5) 0:3:4:5:3 | (5) 2:3:4:5:4 | Digit I of the forelimb formed only by a very reduced metacarpal I; its proximal end is not flat as the other metacarpals, showing a small bulge. Hindlimb follows the pentadactyl condition. |  |
| *Neusticurus bicarinatus* | (5) 2:3:4:5:3 | (5) 2:3:4:5:4 | Both fore and hindlimb follow the pentadactyl condition. |  |
| *Neusticurus rudis* | (5) 2:3:4:5:3 | (5) 2:3:4:5:4 | Both fore and hindlimb follow the pentadactyl condition. Phalangeal formula following Grizante (2009). | Grizante (2009) |
| *Nothobachia ablephara* | dIV | (2) 0:0:2:4:0 | A single digit is present in the forelimb, with one metacarpal and two phalanges, the last one, the ungual phalanx, bearing a nail. The hindlimb is formed by two digits, identified as digits III and IV. |  |
| *Pholidobolus montium* | (5) 2:3:4:5:3 | (5) 2:3:4:5:4 | Both fore and hindlimb follow the pentadactyl condition. |  |
| *Placosoma cordylinum* | (5) 2:3:4:5:3 | (5) 2:3:4:5:4 | Both fore and hindlimb follow the pentadactyl condition. Phalangeal formula following Grizante (2009). | Grizante (2009) |
| *Placosoma glabelum* | (5) 2:3:4:5:3 | (5) 2:3:4:5:4 | Both fore and hindlimb follow the pentadactyl condition. |  |
| *Potamites ecpleopus* (former *Neusticurus ecpleopus*) | (5) 2:3:4:5:3 | (5) 2:3:4:5:4 | Both fore and hindlimb follow the pentadactyl condition. |  |
| *Potamites juruazensis* (former *Neusticurus juruazensis*) | (5) 2:3:4:5:3 | (5) 2:3:4:5:4 | Both fore and hindlimb follow the pentadactyl condition. Phalangeal formula following Grizante (2009). | *Grizante (2009)* |
| *Procellosaurinus tetradactylus* | (5) 0:3:4:5:3 | (5) 2:3:4:5:4 | Osteology of the limbs described in Roscito and Rodrigues (2013). Digit I of the forelimb is represented by a very reduced metacarpal I; hindlimb follows the pentadactyl condition. | *Roscito and Rodrigues (2013)* |
| *Proctoporus bolivianus* | (5) 2:3:4:5:3 | (5) 2:3:4:5:4 | Both fore and hindlimb follow the pentadactyl condition. |  |
| *Proctoporus xestus* (former *Opipeuter xestus*) | (5) 2:3:4:5:3 | (5) 2:3:4:5:4 | Both fore and hindlimb follow the pentadactyl condition. |  |
| *Psilophthalmus paeminosus* | (5) 0:3:4:5:3 | (5) 2:3:4:5:4 | Osteology of the limbs described in Roscito and Rodrigues (2013). Digit I of the forelimb is represented by a very reduced metacarpal I; hindlimb follows the pentadactyl condition. | *Roscito and Rodrigues (2013)* |
| *Ptychoglossus brevifrontalis* | (5) 2:3:4:5:3 | (5) 2:3:4:5:4 | Both fore and hindlimb follow the pentadactyl condition. |  |
| *Rhachisaurus brachylepis* | (5) 0:3:4:5:3 | (5) 1:3:4:5:0 | Digit I in the forelimb and digit V in the hindlimb are represented only by a reduced metacarpal/metatarsal and no phalanges. Kizirian and McDiarmid (1998) agree on the phalangeal formula. | *Kizirian and McDiarmid (1998)* |
| *Scriptosaura catimbau* | Humeurs + distal element | dIV | Osteology of the limbs described in Roscito and Rodrigues (2013). Forelimb absent externally, represented by a rudimentary humerus and a vestigial ossification distal to it. The hindlimb has a single digit, which is formed by one metatarsal and one phalanx which does not bear a nail. | *Roscito and Rodrigues (2013)* |
| *Stenolepis ridleyi* | (5) 2:3:4:5:3 | (5) 2:3:4:5:4 | Both fore and hindlimb follow the pentadactyl condition. The ungual phalanx from digit I of the forelimb is very reduced and does not bear a nail. |  |
| *Tretioscincus agilis* | (5) 2:3:4:5:3 | (5) 2:3:4:5:4 | Both fore and hindlimb follow the pentadactyl condition, although both phalanges from digit I of the forelimb are reduced. |  |
| *Vanzosaura rubricauda* | (5) 0:3:4:5:3 | (5) 2:3:4:5:4 | Osteology of the limbs described in Roscito and Rodrigues (2013). Digit I of the forelimb is represented by a very reduced metacarpal I; hindlimb follows the pentadactyl condition. | *Roscito and Rodrigues (2013)* |

A typical squamate pentadactyl condition (Romer, 1956) is that of 5 digits in both fore and hindlimb, with a specific number of phalanges in each digit (**Figure 2**):

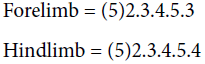

**Figure 2.**
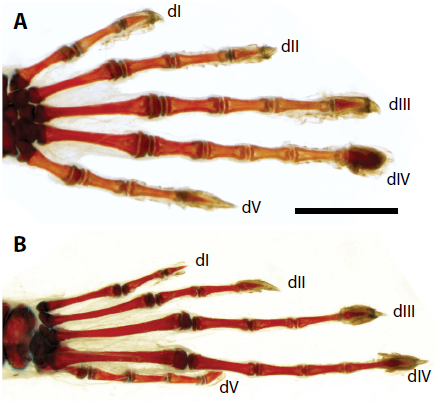
Forelimb (A) and hindlimb (B) of *Cercosaura schreibersi*, showing the generalized phalangeal arrangement. Digit identity is indicated as dI-dV. Anterior to the top. Scale bar = 1mm.

Most gymnophthalmid species have limbs following the generalized pentadactyl arrangement, which is the case for all cercosaurinis and alopoglossinis, and a few other species from the Ecpleopodinae. However, several species that show a full pentadactyl arrangement exhibit a reduction in size of one or both phalanges of digit I of the forelimb; this is seen in at least two gymnophthalmid groups.

The majority of species from the Iphisiini (Rodrigues *et al*., 2009) show a slight reduction in the size of the ungueal phalanx from dI, with *Alexandresaurus, Colobosaura* (**Figure 3D**), and *Stenolepis* (**Figure 3E**) bearing no claw in this digit; in *Iphisa*, both phalanges seem reduced. Reduction is not clear in *Acratosaura*, more material is needed in order to confirm whether its limbs share the morphology of its relatives.

**Figure 3.**
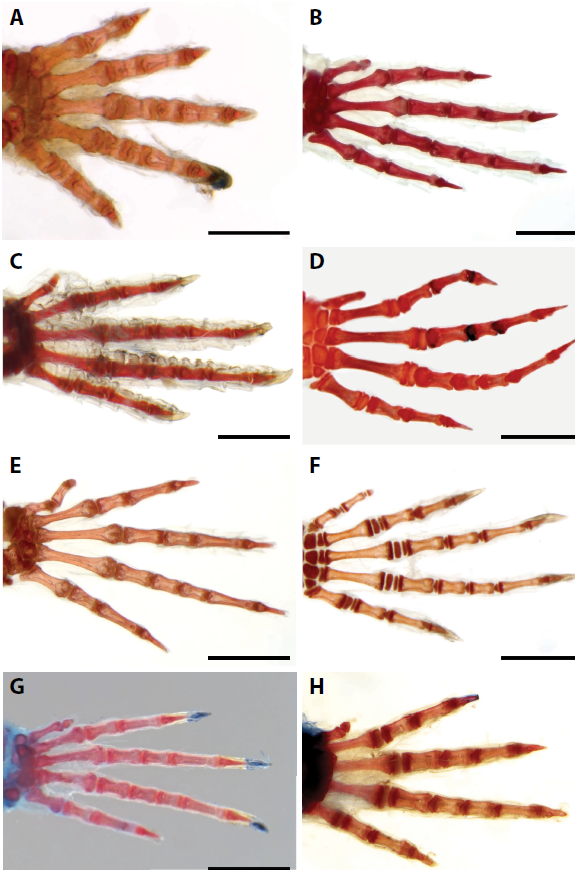
Forelimb morphologies in gymnophthalmid species with minor limb reductions: *Anotosaura vanzolinia* (A); *Colobodactylus dalcyanus* (B); *Colobodactylus taunayi* MTR 00746 (C); *Colobodactylus taunayi* MZUSP 94254 (D); *Colobosaura modesta* (E); *Stenole-pis ridley* (F); *Heterodactylus lundii* (G); *Heterodactylus imbricatus* (H). Digit I to the top. Scale bars: A = 0.5 mm; B-H = 1.0 mm.

*Tretioscincus* is the other genus in which a clear reduction in size of phalanges in dI can be observed; the other gym-nophthalmini species have lost either one *(Gymnophthalmus*) or both phalanges from d1 of the forelimb (*Micrablepharus, Psilophthalmus, Procellosaurinus, Vanzosaura)* (**Figure 4 A-D**).

**Figure 4.**
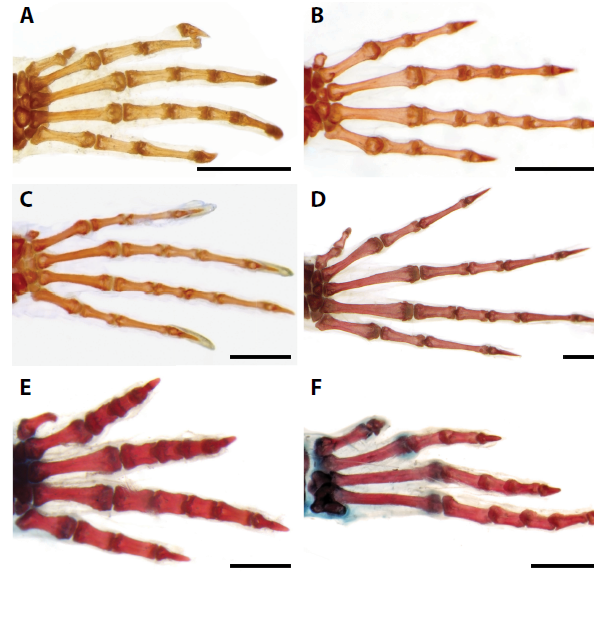
Limb morphologies in gymnophthalmid species with minor limb reductions: *Gymnophthalmus underwoodi* forelimb (A); *Micrablepharus atticolus* (B); *Psilophthalmus paeminosus* (C); *Tretioscincus agilis* (D); *Rachisaurus brachylepis* (E, F). Digit I to the top. Scale bars: A, B, D, F = 1.0 mm; C, E = 0.5 mm

Complete loss of dI of the forelimb, as seen in the beforementioned gymnophthalmini species, only occurs in *Rachisaurus brachylepis* (**Figure 4 E,F**). *Colobodactylus*’ dI of the forelimb is clearly reduced, but we disagree with Kizirian and McDiarmid (1998) and Grizante (2009), who report only the presence of mcI, since we have observed one reduced phalanx following the metacarpal (mc) in both *C. dalcyanus* and *C. taunayi*. However, it is possible that there is intrapopulational variation for this character; the presence of a vestigial ossification distal to mc1 in *C. taunay* (MTR 746) and the apparent absence of such element in the second individual analysed here (MZUSP 94254) argues in favor of this variation (**Figure 3B,C**).

Interestingly, all those species in which both phalanges from dI of the forelimb are lost, the corresponding metacarpal (mcI) is always present. Also worth of notice is that in all these cases previously mentioned, the minor reductions, or loss of dI, are not accompanied by reductions/losses in the hindlimbs, which maintain the pentadactyl arrangement. The exception is *Rachisaurus brachylepis*, which has lost one phalanx from dI, and all phalanges from dV of the hindlimb; this condition is not seen in any of the other Gymnophthalmidae analysed.

Another “type” of reduction is seen in some species of the Ecpleopodinae: while most are pentadactylous, *Anotosaura* and *Dryadosaura* exhibit a loss of the last phalanx of dIV of the forelimb. In addition, in *Anotosaura collaris* the last phalanx of dV of the hindlimb is absent (**Figure 3A**; its sister species, *A. vanzolinia*, shows the full pentadactyl arrangement). The single individual of *Colobosauroides cearensis* analysed shows a left-right asymmetry in the number of phalanges in dIV of the forelimb.

Limb reduction reaches extreme cases in two gymnoph-thalmid lineages: the gymnophthalmini, with *Nothobachia* and *Calyptommatus* (and possibly *Scriptosaura;* Rodrigues and dos Santos, 2008), and the Bachiini (**Figure 5**).

**Figure 5.**
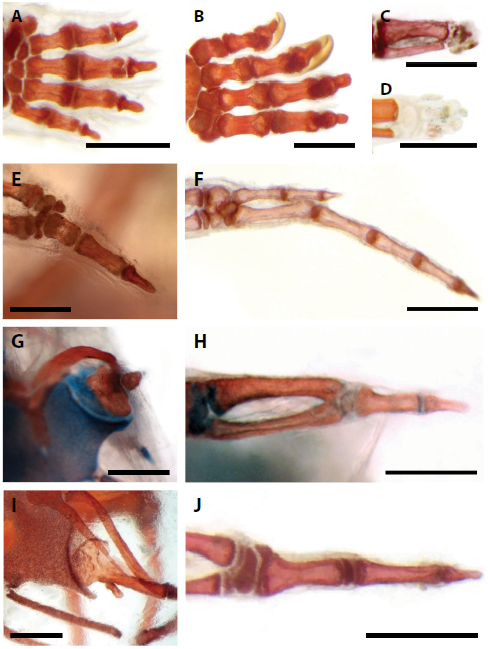
Limb morphologies in gymnophthalmid species with the greatest degrees of limb reduction: *Bachia panoplia* fore (A) and hindlimb (B); *Bachia bresslaui* hindlimb (C); *Bachia flavescens* hindlimb (D); *Nothobachia ablephara* fore (E) and hindlimb (F); *Scriptosaura catimbau* fore (G) and hindlimb (H); *Calyptommatus leiolepis* forelimb (I); *Calyptommatus sinebrachiatus* hindlimb (J). Digit I to the top. Scale bars: A-C, F, G, I-J = 0.5 mm; D, E, H = 0.25 mm.

Within the Gymnophthamini (**Figure 5E-J**), *Nothobachia* still retains both fore and hindlimbs, although they are very reduced (forelimb styliform and hindlimb with two digits) and presumably not functional for fast walking (Renous *et al*., 1998). *Scriptosaura* and *Calyptommatus* show even further reductions: both lack the forelimb, although retaining a vestigial internal humerus close to the pectoral girdle (and possibly, a vestige of the second limb segment in *Scriptosaura;* Roscito and Rodrigues, 2013); both species have a styliform hindlimb.

*Bachia* is a very interesting genus with respect to the diversity of limb morphologies. Limb reduction within the group is more pronounced in the hindlimbs than in the forelimbs, the opposite of all the other gymnophthalmid species in which some kind of reduction occurs. More interestingly, the digits of *Bachia* species show an apparent loss of anterior-posterior patterning (**Figure 5A,B**), as previously noted (Kohlsdorf and Wagner, 2006). Digit identity can be determined in species with more developed limbs, but in cases of extreme reductions, no recognizable digit structure is left (**Figure 5C**).

*Bachia* species are quite rare in the field, and also in museum collections, therefore, we could only get access to three cleared and stained specimens; the majority of information for these species was compiled from the literature. Because these are rare animals, the taxonomic studies on *Bachia* species usually don’t contain osteological information, and descriptions of the limbs are mainly based on external morphology. For this reason, we relied solely on the osteological information published by Dixon (1973), Thomas (1965), Kizirian and McDiarmid (1998), and Kohlsdorf and Wagner (2006).

### Phylogenetic analysis

The types of limb reduction described above cluster in specific groups (**Figure 6 and Figure 7**): in the lineage that holds *Rachisaurus*, heterodactylinis, iphisiinis, and gymnophthalminis, reduction affects digit I of the forelimb, while the hindlimb is frequently pentadactylous (except for *Rachisaurus brachylepis). Bachia* species exhibit a wide range of morphologies, but none of them following the typical pentadactyl arrangement. Last, reduction in the ecpleopodinis affects digit IV of the forelimb and also digit V of the hindlimb.

**Figure 6.**
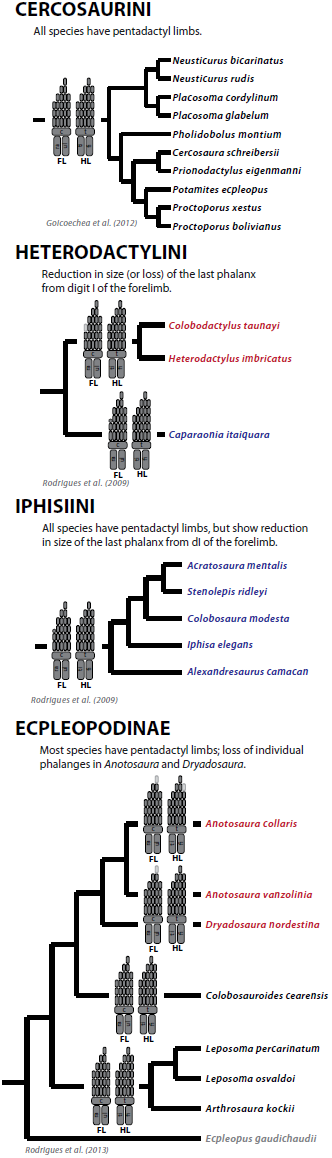
Evolutionary relationships within the Cercosaurini, Heterodactylini, Iphiisini, and Ecpleopodinae subgroups, with a representation of the different limb morphologies observed in each species/groups of species. Limb skeletal elements present in the limbs are colored dark grey, and those that are absent (in reference to the generalized pentadactyl condition) are colored light grey. Species names colored in black indicate those species that have fully pentadactyl limbs; names colored in blue indicate those species that also have pentadacatyl limbs but show reduction in size of one or more phalanges; species colored in red have lost one or more phalanges in relation to the pentadactyl condition. FL/HL, forelimb/hindlimb; ra, radius; ul, ulna; c, carpus; ti, tibia; fi, fibula; t, tarsus.

**Figure 7.**
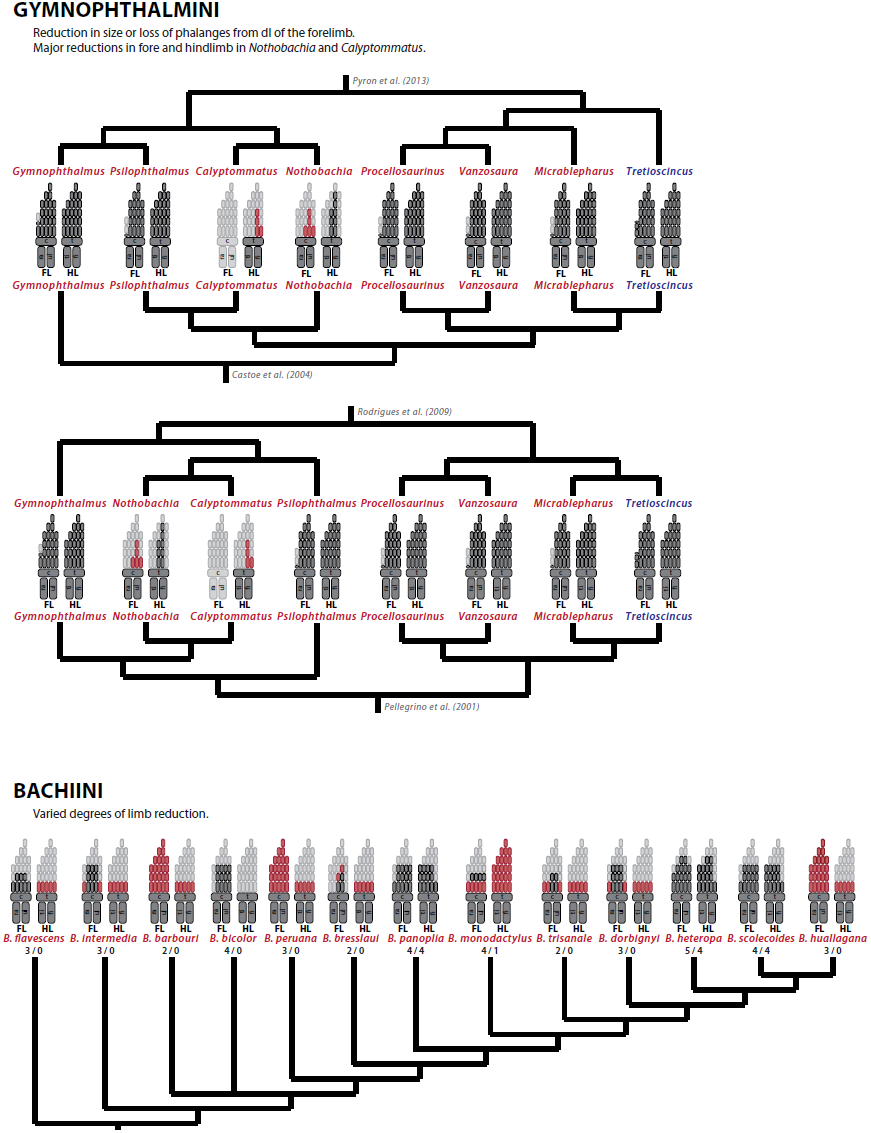
Evolutionary relationships within the Gymnophthalmini and Bachiini subgroups, with a representation of the different limb morphologies observed in each species. Limb skeletal elements present in the limbs are colored dark grey, those that are absent (in reference to the generalized pentadactyl condition) are colored light grey, and those with uncertain identity are in red. Species names colored in blue indicate those species that have pentadacatyl limbs but show reduction in size of one or more phalanges; species colored in red have lost one or more phalanges in relation to the pentadactyl condition. FL/HL, forelimb/hindlimb; ra, radius; ul, ulna; c, carpus; ti, tibia; fi, fibula; t, tarsus.

Mapping the distribution of limb morphologies onto the gymnophthalmid subgroups shows shared morphologies among groups of species and allows pinpointing shared or independent origins for similar limb arrangements.

In general, the species that show some kind of limb reduction are grouped together. For example, limb reduction in the Ecpleopodinae is restricted to the genera *Anotosaura* and *Dryadosaura*, which are nested within the group, while all sister species show no kind of reduction (**Figure 6**). The same is true for the Heterodactylini, with the reduced forms grouping together (**Figure 6**). Cercosaurini and Iphisiini species don’t exhibit variation in limb morphology; the former is comprised of only pentadactyl species, while species from the later group also share the typical pentadactyl arrangement, but show reduction in size of the last phalanx of digit I of the forelimb (**Figure 6**).

Gymnophthalmini and Bachiini, on the other hand, do not show a clear clustering of species with shared phenotypes (**Figure 7**). One of the two main branches recognized among the Gymnophthalmini (**Figure 7**) is comprised exclusively of the lacertiform *Tretioscincus, Micrablepharus, Vanzosaura*, and *Procellosaurinus;* while the other holds the lacertiform *Gymnophthalmus* and *Psilophthalmus*, and the snake-like *Nothobachia* and *Calyptommatus (Gymnophthalmus* is placed as a separate lineage by Castoe *et al*., 2004). Loss of digit I of the forelimb occurs in *Psilophthalmus, Micrablepharus, Vanzosaura*, and *Procellosaurinus*, yet *Psilophthalmus* does not group directly with the later three species. The snake-like gymnophthalminis *Nothobachia* and *Calyptommatus*, which represent the extremes of this subgroup regarding limb morphology, are always grouped together (the topology of Castoe *et al*., 2004, however, places *Psilophthalmus* in between them). The Bachiini (**Figure 7**) show an even more puzzling situation, in which the diversity of limb configurations follows no clear pattern among the species for which limb morphology is known in detail. The limiting amount of data available regarding both the anatomy and evolutionary relationships of *Bachia* species contribute to this confusing scenario.

### Analysis of gene expression in *Calyptommatus sinebrachiatus*

We analysed the expression of Shh in the fore and hindlimb buds of *Calyptommatus sinebrachiatus* embryos staged from 6 to 16 days of development (**Figure 8**). When the egg is laid, both fore and hindlimb are present (Roscito and Rodrigues, 2012b); our selection of developmental stages was intended to sample the early development of both limbs. The forelimb bud develops up to 9-10 days, and then its growth stops and the bud starts to degenerate. The hindlimb bud, which does not degenerate, is already paddle-shaped at 16 days after laying, with femur, tibia, and fibula detected by cartilage staining (Roscito and Rodrigues, 2012a).

**Figure 8.**
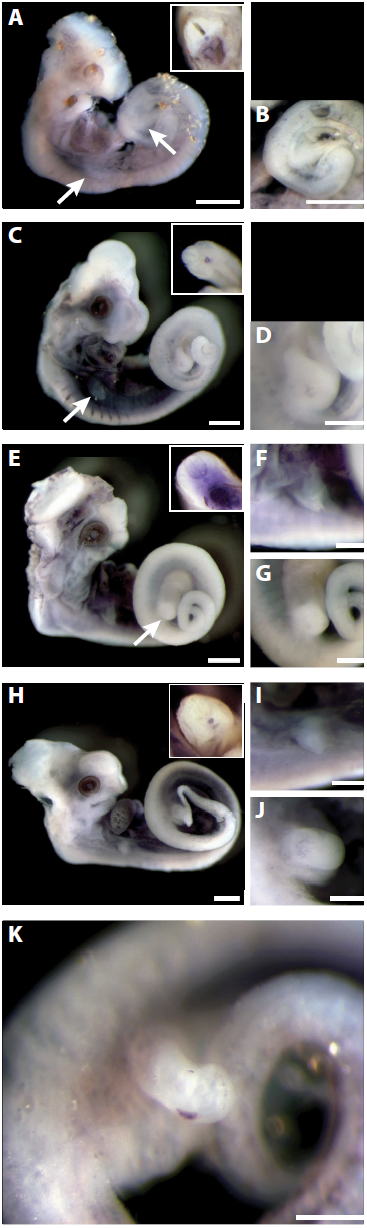
Expression of the gene Sonic Hedgehog (Shh) in embryos of the gymnophthalmini *Calyptommatus sinebrachiatus* with 6 (A-B), 10 (C-D), 12 (E-G), 14 (H-J), 16 (K) days of development. Images A, C, E, and H show the entire embryo: arrows in A point to the fore and hindlimb, arrow in C points to the forelimb, and arrow in E points to the hindlimb; the images in detail on the upper right corner represent a transversal cut at mid-trunk showing positive staining in the nothocord. Forelimbs are represented in F and I; hindlimbs in D, G, J, and K. Scale bars: A-B, C, E, H = 0.5mm; D, F, G, I, J, K= 0.25mm.

Sonic hedgehog mRNA was consistently observed in the notochord of all embryos (**Figure 8A, C, E, H; images in detail**). However, it was not detected in the forelimb at any of the stages analysed. In contrast, Shh it was detected in the posterior mesenchyme of the hindlimb bud of the 16-day old embryo (**Figure 8K**), the oldest stage analysed.

## DISCUSSION

In addition to the knowledge of the molecular pathways involved in ‘normal’ limb development, there is also a vast amount of information on the influence of disruptions of such pathways to the resulting limb morphology. These interventions, and the phenotypes resulting from it, provide starting points for investigations of limb development in other species that show different limb morphologies.

### Patterns of limb reduction in the Gymnophthalmidae

Limb reduction has evolved repeatedly in squamates, with almost every major group showing the loss of one or more bones in the limbs. The situation is no different among the Gymnophthalmidae, with all but three groups exhibiting loss of one or more phalanges (**Figure 1**).

Minor reductions, involving the loss of one or two phalanges, account for the majority of cases of limb reduction among the gymnophthalmids. Digit I of the forelimb is the most affected: out of the 10 genera analysed that show these minor losses, 8 have lost phalanges in dI. When a single phalanx is lost, the remaining phalanx is very reduced in size, while the size of the metacarpal seems comparable to that of the other metacarpals. In contrast, when both phalanges are absent, mcI is reduced to a vestigial element. Phalanx loss is also seen in dIV of the forelimb, but with a lower frequency when compared to losses in dI. This scenario contrasts the observations of Greer (1991), that has shown that dIV is more likely to be affected when losses occur in a single digit from the forelimb (26 times), followed by dI (6 times), and last, by dV (only 1 case).

Losses of phalanges in either dI or dIV of the forelimb seem to be specific to distinct lineages: loss in dI is seen in the Rachisaurinae, Heterodactylini, Iphisiini and Gymnophthalmini, while loss of dIV occurs only in some species of the Ecpleopodinae. However, Rodrigues *et al*. (2013) have recently described a new *Leposoma* species, *L. sinepollex* that, as the name suggests, lacks dI of the forelimb. Furthermore, the closely-related *L. nanodactylus* is reported to have a shorter dI of the forelimb (Rodrigues 1997; Rodrigues *et al*., 2013). These observations break down the notion of clade-specific types of reduction, since *Leposoma* are nested within the Ecpleopodinae, a group characterized by reductions affecting dIV of the forelimb. Furthermore, it shows that the *scincoides* lineage of the *Leposoma* genus, to which *L. nanodactylus* and *L. sinepollex* are allocated, represents one more independent instance of digit reduction among the gymnophthalmids (Rodrigues *et al*., 2013). A second event of loss of dI of the forelimb among the Ecpleopodinae is seen in the monotypic *Amapasaurus tetradactylus* (Cunha, 1970; pers. obs.). This species seems to be more related to the *parietale* group of *Leposoma* (Rodrigues and Ávila-Pires, 2005), which holds the two *Leposoma* species analysed here (*L. percarinatum* and *L. osvaldoi*), none of which showing any kind of limb reduction. If this relationship is confirmed, then *Amapasaurus* represents yet another independent event of limb reduction within the group.

Interestingly, the hindlimbs of all of those gymnophthalmid species that show minor losses of phalanges in the forelimb do not show any kind of reduction, and still maintain the ancestral phalangeal formula. The exceptions are *Anotosaura collaris*, which has lost a single phalanx from dV, and *Rachisaurus brachylepis*, which has lost the entire dV but the corresponding metatarsal. Digit V of the hindlimb is the digit that accounts for the most cases of losses of phalanges (Greer, 1991; Shapiro *et al*., 2007), although losses of all phalanges from dV usually co-occur with complete loss of dI (Shapiro *et al*., 2007), which is not the case for *Rachisaurus*.

The effect of intrapopulational variation on the number of phalanges lost in both fore and hindlimb, as previously observed in *Bachia* and *Hemiergis* (Dixon, 1973; Choquenot and Greer, 1989), cannot be estimated based on our sampling, most of which consisting of a single individual per species. The observation of two specimens of *Colobodactylus taunayi* with a different configuration of dI of the forelimb - depending on the interpretation of the vestigial ossification seen in specimen MZSUP 94254 (**Figure 3D**), specimens either differ in the number of phalanges, or in the morphology of the single phalanx - argues in favor of the need increased sampling in order to understand the morphology of reduced limbs and its dominant pattern within a species. On the other hand, we have analysed several individuals of *Vanzosaura rubricauda, Psilophthalmus paeminosus*, and *Procellosaurinus tetradactylus* and none of those showed any variation in number of phalanges. Interestingly, the three later species have a vestigial mcI, while that of *Colobodactylus* has a normal size. The extent to which this variability in the number of phalanges represents a clade-specific flexibility, some kind of developmental constraint related to how reduced is the digit, or a simple factor of change in our sampling, will unfortunately remain unknown until increased sampling and more detailed anatomical studies are done.

Major limb reductions are seen in two other gymnophthalmid groups: the Gymnophthalmini, with *Nothobachia, Calyptommatus*, and *Scriptosaura* (Rodrigues and dos Santos, 2008), and the genus *Bachia*. In such severe cases of limb reduction, especially regarding losses of multiple digits and of other elements to an almost limblessness state, the remaining skeletal elements (if any) are vestigial and frequently show some kind of loss of anatomical information. Examples of limb elements that don’t share anatomical similarity to its homologous elements from pentadactyl relatives are: the vestigial humerus of *Calyptommatus* and *Scriptosaura;* the extremely reduced elements seen laterally to the digits in the forelimb of *Nothobachia* and in the hindlimb of *Calyptommatus*, which cannot be identified only based on the adult morphology (**Figure 5A, J**); and the vestigial elements in *Bachia bresslaui* hindlimb (**Figure 5C, D**).

On the other hand, the skeletal elements present in “intermediate forms” usually show some degree of anatomical individuality and may still maintain topological relations to neighboring elements. This is the case for the hindlimb of *Nothobachia ablephara*, in which the two remaining digits can be identified as dIII and dIV based on their relations to the tarsal elements (Roscito and Rodrigues, 2013). Remarkably, digits and phalanges in *Bachia* seem to be deprived of anatomical information, in the sense that all digits - even in species with high number of digits, look similar to each other.

Two major patterns of limb reduction can be recognized among those Gymnophthalmid lizards that show some degree of limb reduction, whether it is a loss of a single phalanx or the loss of essentially the entire limb. In all species but *Bachia*, reduction of the forelimb is always more advanced than reduction in the hindlimb; *Bachia* shows an opposite trend. The first case, in which the forelimb is more reduced than the hindlimb, is the most frequent pattern in limb-reduced squamates, while the second case is only seen in *Bachia*, in one scincid and one teiid, and in the amphisbaenian genus *Bipes* (Brandley *et al*., 2008). As every ‘rule’ has its exception, two recently described *Bachia* species (*B. micromela* and *B. psamophila*; Rodrigues *et al*., 2007) have the forelimbs more reduced than the hindlimbs. Although no osteological data was presented for either species, the representations clearly show that the hindlimb is more developed than the forelimb (both in length and in number of digits, in the case of *B. psamophila)*.

In addition, reduction in the fore and hindlimbs of *Bachia* seem to follow distinct morphological patterns: reduction in the forelimb has, apparently, an anterior predominance (dI usually reduced or absent), while in the hindlimb, dV is the most affected. Reduction/loss of dI of the forelimb is common among limb-reduced forms (75% of the cases; Shapiro *et al*., 2007), and there is not a correlation between reduction/loss of this digit with reduction/loss of any of the other digits, meaning di is relatively independent. Although reduction in dV of the hindlimb is also common (90% of the cases; Shapiro et al., 2007), it is highly correlated with corresponding reduction in dI of the hindlimb (73%). Digit V is lost in the hindlimbs of *B. panoplia, B. scolecoides*, and *B. heteropa*, but dI is complete (2 phalanges) in all three species, contrary to the co-loss of dI and dV reported in the above-mentioned study. This predominant reduction of the post-axial side of the limb, with maintenance of the pre-axial side, resembles the pattern seen in archosaurs fore and hindlimbs (Shapiro *et al*., 2007) and in anurans (Shubin and Alberch, 1986). In addition, a few *Bachia* species such as *B. panoplia, B. scolecoides, B. intermedia, B. bicolor*, and *B. dorbignyi*, show morphologically similar digits with uniform phalangeal numbers, similar to what is seen among the turtles and in some archosaurs (Shapiro et al., 2007). These opposing modes of digit reduction between fore and hindlimbs of *Bachia* suggest that different developmental mechanisms may be coming into play. Unfortunately, the lack of a detailed anatomical analysis of the genus is a drawback to further hypothesis on limb development.

### Developmental biology of reduced limbs

#### Morphological perspective

The development of the tetrapod limb is a complex process that requires precise special and temporal coordination of many signaling molecules that lead to the differentiation of an initially homogeneous population of cells into the different tissues that form the limbs.

One of those tissues, the limb skeleton, arises from pre-cartilaginous primordial that are formed in a proximal-distal direction. In amniotes, the order of appearance of limb skeletal elements seems to be remarkably conserved, despite the major differences in limb morphology across species. Comparative analyses have shown that the pre-cartilaginous elements form following a primary axis of development that runs through the humerus/femur, ulna/ fibula, distal carpal/tarsal IV, and digit IV; the remaining digits usually form following the sequence III > II/V > I (Shubin and Alberch, 1986). This generality holds true for the squamates analysed so far (Howes and Swinnerton, 1901; Mathur and Goel, 1976; Rieppel, 1994; Shapiro, 2002; etc).

Another generality derived from the analysis of limb-reduced species is that limb elements are lost in the reverse order of their development; thus, the last elements to form are usually lost first. In the case of digits, loss follows the order I > V/II > III/IV (Morse, 1872; Lande, 1978; Greer, 1987; 1991; Shapiro *et al*., 2007). This scenario could imply that losses of elements that are not essential for the development of other elements could take place more easily than loss of those elements that form the primary axis of the limb. The apparent constraint against the loss of the central digits (III and IV) could also be related to the role that these digits play in hand/foot stability during locomotion (Greer, 1991), but we still lack a detailed and comparative knowledge of how hand/feet width and length could influence on locomotion performance.

Taking into account the prevalence of the primary axis, and the relative common sequence of digit development, one could preliminarily infer that the reduced digits of *Calyptommatus, Scriptosaura*, and *Nothobachia* are the most central digits: those in *Nothobachia* hindlimb would be identified as dIII and dIV, and the single digit in *Nothobachia* forelimb and in *Calyptommatus* and *Scriptosaura* hindlimbs would be identified as dIV. However, if identified as such, then the number of phalanges in these digits does not match to that observed in the corresponding digit if a generalized pentadatyl limb.

The first question that stems from this observation is if these apparently incomplete digits would be the product of an early termination of limb bud development (see Shapiro, 2002, for further discussion). If so, we should expect to find a similar digit/phalangeal configuration in some developmental stage of a pentadactyl limb bud. However, this is not the case for neither species, arguing against a truncation mechanism to explain the resulting limb morphology.

In contrast, there are cases in which the adult phalangeal arrangement corresponds to the arrangement seen in a particular developmental stage, thus reflecting that the reduction process was likely the result of a truncation of embryonic development. This is the case for *Hemiergis initialis* (Shapiro, 2002), and several other skinks, agamids, cordylids, gekkonids, and others, that show minor losses of one or two terminal phalanges of digits IV and V (Greer, 1991; Russel and Bauer, 2008); among the gymnophthalmids, *Anotosaura* shows a phalangeal configuration resembling that of a late developmental stage of *Calotes versicolor* (Mathur and Goel, 1976). Interestingly, all these cases involve minor losses of phalanges (1-3 phalanges lost).

Digit configurations seen in some *Bachia* species, such as B. *panoplia*, *B. scolecoides*, and *B. bicolor* also do not seem to correspond to truncations of a pentadactyl embryonic developmental program. On the other hand, the forelimbs of *B. bresslaui, B. trisanale*, and *B. heteropa* resemble stages of development of other squamates (Shapiro *et al*., 2007), and could possibly have originated from truncations of development.

#### Molecular signaling perspective

The understanding of the signaling network that controls limb growth and patterning during embryonic development sheds light on the possible mechanisms by which a limb becomes reduced. Lab-induced mutations, or alterations of individual signaling pathways, result in a variety of limb phenotypes, some of which may resemble those seen in natural populations. Furthermore, the improvement and easier access to genetic and molecular tools led to an increasing interest in the investigation of a wider diversity of species other than the commonly studied chicken and mouse. This increasing knowledge helps to direct investigations on species never so far studied.

Multiple signals interact to control both the growth and patterning of the developing limb. Changes in such system can affect growth, but not the overall pattern (scaling), or can affect the pattern itself (and, hence, localized growth) and result in a different configuration of the skeletal elements.

The forelimb of *Dryadosaura* looks like the result of a scaling process of limb autopod, since the phalanges are much shorter in comparison to those of the hindlimb, and of the forelimbs of similar-sized gymnophthalmids. However, *Dryadosaura* also has one phalanx less from the generalized pentadactyl pattern - a condition seen in many other Gymnophthalmids. These frequent asymmetric losses of phalanges reflect some kind of deviation from the common developmental pattern other than a simple scaling mechanism, which should, in principle, affect all digits equally.

Digits form from single cartilaginous condensations in the autopod, which are separated from each other by interdigital spaces. These condensations elongate and segment into phalanges, and the distal-most phalanx differentiates into the ungueal phalanx, associated with epidermal structures. Growth of the digit cartilaginous condensation is controlled by FGF8 signaling from the apical ectodermal ridge (AER), which, in conjunction with WNT signaling from the ectoderm, maintain the distal-most cells in an undifferentiated, proliferative state (Stricker and Mundlos, 2011). BMPs from the mesenchyme are important positive regulators of chondrogenesis. Downregulation in either FGF8 or BMP signaling results in shorter cartilaginous condensations and, frequently, in brachydactyly phenotypes (reduced size or loss of phalanges; Sanz-Esquerro and Tickle, 2003; Stricker and Mundlos, 2011). Conversely, over expression of FGF8 results in increased growth of the digit condensations, and in the formation of additional phalanges in the chicken limb (Sanz-Esquerro and Tickle, 2003).

In this scenario, minor spatial and temporal modulations of FGF8 or BMPs expression could account for the formation of a reduced number of phalanges, or the reduction in size of these elements in the gymnophthalmids analyzed here and also in other amniotes with reduced limbs. Furthermore, the predominance of loss/reduction in the lateral-most digits (dI and dV), a common trend among squamate reptiles (Greer, 1991; Shapiro *et al*., 2007), could be easily explained by a possible antero-posterior shortening of the extension of the AER, which would put the lateral-most digits under the influence of smaller amounts of FGF signaling. In fact, shortening of the AER is likely one of the mechanisms that contribute to the reduction of dII and dV in the embryonic limbs of some mammals (Cooper *et al*., 2014; Lopez-Rios *et al*., 2014).

In contrast, the more infrequent cases of loss of the terminal phalanx from digit IV (seen here only in *Anotosaura* species) could be explained by an early termination of AER-derived signaling, which would lead to the truncation of the normal developmental sequence. This hypothesis is supported by experimental manipulations in chicken limbs, in which early interruption of FGF signaling result in the loss of terminal phalanges and in the premature induction of the digit tip developmental program (Sanz-Esquerro and Tickle, 2003) - in fact, the modulation between lower FGF signaling and the switching on of the “tip developmental program”, which involves Wnt signaling, explains the presence of the ungueal phalanx in those digits with reduced number of phalanges. Furthermore, the ungueal phalanx from dIV is the last one to form in development (Mathur and Goel, 1976; Shapiro *et al*., 2007), hence, would be the first one lost following an early termination of FGF signaling.

Signaling from the AER promotes mesenchymal cell proliferation and limb outgrowth; inhibition of FGF8 and FGF4 at different time developmental stages leads to progressive loss of elements along the proximo-distal axis, and knockdown of these FGFs results in failure of limb formation (Lewandoski *et al*., 2000; Sun *et al*., 2002; Dudley *et al*., 2002). Downregulation of AER signaling results in reduced cell proliferation, resulting in skeletal defects due to the reduced number of cell progenitors. Raynaud (1990) showed that inhibition of DNA synthesis results in a reduction of the number of digits formed in the limbs of the pentadactylous lizard *Lacerta viridis*, and that the earlier the treatment, the more digits we missing. More interestingly, the digits most frequently affected (lost) were dI and dV and those most frequently retained were dIII and dIV, which corresponds to the naturally occurring pattern described previously (Greer, 1991; Shapiro *et al*., 2007).

A premature downregulation of FGFs from the AER and the consequent truncation of the developmental pathway could also explain the morphology of the forelimb of *Bachia heteropa*, which has a phalangeal arrangement (1.2.3.3.2) similar to that seen in a developmental stage of *Hermiergis initialis* (Shapiro, 2002). However, the arrangement of *B. heteropa* hindlimb (2.2.3.3.0) is not paralleled by any developmental stage of any squamate studied so far, although it is similar to a developmental stage of the newt *Ambystoma* (Alberch and Gale, 1985).

The uniform phalangeal number seen in most *Bachia* species (1-2 phalanges in at least 3 digits) could be the result of truncations of the developmental program coupled with additional mechanisms directing the loss of lateralmost digits - as for example the already-mentioned antero-posterior shortening of the AER and of the signaling derived from it.

The homogeneity of both size and morphology of the digits of those *Bachia* species led Kohlsdorf and Wagner (2006) to suggest a correspondence of these arrangements with the phenotype resulting from mouse knockout mutants for Gli3. This mesenchymal transcription factor, together with Sonic hedgehog from the ZPA (zone of polarizing activity), are key players in the antero-posterior polarization of the developing limb bud (Wang *et al*., 2000; Litingtung *et al*., 2002; te Welscher *et al*., 2002); their antagonistic interactions determine a signaling gradient along the AP axis of the limb bud over which digit number and identity are laid down (Litingtung *et al*., 2002). Limbs that develop in the absence of Gli3 are polydactylous, and the resulting digits have 2 phalanges each and seem identical (“appear more serially homologous than in the wild type”; Litingtung *et al*., 2002). Although we cannot be sure about the developmental mechanisms responsible for the limb phenotypes seen in *Bachia*, it sounds unlikely that some sort of downregulation of Gli3 could account for these arrangements. Reduction in the amount of Gli3 would imply an anterior expansion of Shh expression, which would result, most likely, in some degree of polydactyly.

Loss or downregulation of Shh, on the other hand, could explain the limb morphologies of *Nothobachia* and *Calyptommatus*. Progressively inactivation of Shh expression from mouse limb buds leads to a corresponding progressive loss of digits (Zhu *et al*., 2008), with more digits being lost the earlier Shh is inactivated. The same effect was observed in scincid lizards from the genus *Hemiergis* with different limb morphologies: reduction in the number of digits is to a reduction in the duration of Shh expression; the more reduced the limb, the less time Shh is present in the limb buds (Shapiro *et al*., 2003). Shh maintains a positive feedback loop with growth-promoting FGFs from the AER via Gremlin signaling (Laufer *et al*., 1994; Niswander *et al*., 1994; Zuniga *et al*., 1999; Lewandoski *et al*., 2000; Sun *et al*., 2002); thus, decrease in Shh expression is always correlated with a decrease in cell proliferation, resulting in a smaller limb bud (Chiang *et al*., 2001; Shapiro *et al*., 2003) and, consequently, in less progenitor cells.

Determining the identity of digits present in partially or greatly reduced limbs can be quite difficult. Even though one might rely on the primary axis generalization, reduced limbs/digits can show a high degree of anatomical divergence from the ancestral condition, often resulting in controversial homology assignments; one of the greatest examples is the over 150-year old debate over the identity of the digits in the chicken wing (reviewed in Young *et al*., 2011). Therefore, integrating morphological and molecular evidences is essential for the elaboration of an evolutionary scenario to accommodate the divergent observations (for example, Young *et al*., 2009).

As discussed above, from the morphological point of view, multiple embryological evidences (Shubin and Alberch, 1986; Müller and Alberch, 1990; Chiang *et al*., 2001; Noro *et al*., 2009; Young *et al*., 2009; Leal *et al*., 2010; Shapiro, 2002, and many others) show that: i) the condensation corresponding to dIV is the first one to form in the limb buds; ii) digits most often form sequentially in a posterior-anterior order (IV>III>II/V>I; although Zhu *et al*., 2008 evidences claim for more detailed observations of digit development); and iii) that digits are lost in the reverse order of their development, which makes dIV the last one to disappear (Morse, 1872; Sewertzoff, 1931; Chiang *et al*., 2001; Shapiro *et al*., 2002).

From the molecular biology point of view, there are still standing questions over the exact roles of Shh signaling in the patterning of the AP axis and in determining digit number and identity (Tabin and McMahon, 2008; Harfe, 2011). Nevertheless, it is clear that, in the absence of Shh signaling, the single digit that forms is the Shh-independent dI (Chiang *et al*., 2001; Ros *et al*., 2003), whereas progressive loss of Shh leads to dIV being lost last (Zhu *et al*., 2008). The detection of Shh in the hindlimb of *Calyptommatus* is an indication that the single-digit phenotype is different from the mutant single-digit phenotype. In addition, Shh-null mice also show severe defects in the zeugopod (Chiang *et al*., 2001), not seen in *Calyptommatus* hindlimb. By combining molecular and morphological inferences, we could suggest that dIV is the remaining digit in the vestigial hindlimb of *Calyptommatus* (as previously interpreted in Roscito and Rodrigues, 2013), and that the absence of other digits may be explained by a failure in maintenance of Shh expression, as observed in *Hemiergis* lizards.

On the other hand, positional information may not always be a good proxy for determining digit identity and homology, given that patterning of digit identity can be uncoupled from the anatomical positioning of the cartilage condensation. The digits in the chicken wing and in both fore and hindlimb of the lizard *Chalcides chalcides* are examples where this uncoupling occurs: while anatomical analysis supports the identification of the digits as dI, dII, and dIII (in both the chicken and *C. chalcides*), gene expression analyses during limb development show that the digits are patterned as dII, dIII, and dIV (Wagner and Gauthier, 1999; Young *et al*., 2009). The Frame Shift Hypothesis (Wagner and Gauthier, 1999) accommodated these conflicting evidences by proposing a homeotic transformation of character identity.

The phalangeal number in the single digit of *Calyptommatus* hindlimb is reminiscent of dI from a pentadactyl condition, which might reflect a frame shift-like mechanism taking place in the patterning of this digit. Digit i develops in the anterior-most area of the hand/foot autopod where only Hoxd13, out of the distal Hoxd genes, is expressed (Montavon *et al*., 2008); thus, the absence of other distal Hoxd genes from the presumptive dI region is a reliable indication of di fate - in fact, Hoxd11 and Hoxd12 expression have been used to identify the homeotic frame shifts of the chicken and the lizard *C. chalcides*, as previously discussed. The investigation of the expression of the distal Hoxd genes in the hindlimb of *Calyptommatus* would be decisive in determining if a homeotic shift would explain the digit configuration.

The extent to which disruptions in Shh signaling may be associated with changes in digit identity is still unknown. On one side, a change in downstream Shh signaling in bovine embryos results in a medial-distal shift of Hoxd13 expression, but the resulting digits do not have dI identity (Lopez-Rios *et al*., 2014). On the other hand, a shift in digit identity was observed in experimentally induced inhibitions of Shh in the chicken wing: digits I and II developed, but these digits formed in embryonic positions of dIII and dIV (Vargas and Wagner, 2009). Therefore, even though Shh was observed in *Calyptommatus* hindlimb, we cannot exclude the possibility of a downstream effect on its signaling cascade that could potentially induce a frame shift event.

Whether similars mechanisms take place in the single-digit fore and hindlimb of *Nothobachia* and *Scriptosaura*, respectively, remains an open question.

The even more reduced forelimb of *Calyptommatus*, represented in by a vestigial humerus located within the body wall, forms normally in early development but regresses at later stages (Roscito and Rodrigues, 2012b). Sonic hedgehog was not detected in this limb bud at any stage, showing that: i) Shh is not necessary for the emergence of the limb bud (consistent with previous results; Chiang *et al*., 2001; Ros *et al*., 2003); and ii) the absence of this important patterning signal may be one of the factors involved in degeneration of the forelimb bud.

Arrested development of limb buds in limb-reduced lizards has been documented previously for the lizards *Anguis fragilis, Ophisaurus apodus*, and *Scelotes brevipes* (Rahmani, 1974; Raynaud, 1963; Raynaud *et al*., 1975); the python snake has vestigial limb buds that do not degenerate (Cohn and Tickle, 1999; Boughner *et al*., 2007). Cetaceans hindlimb buds also form during embryonic development but degenerate (Sedmera *et al*., 1997; Thewissen *et al*., 2006). Remarkably, Shh is not expressed in the hindlimbs of both the dolphin and snake (Thewissen *et al*., 2006; Cohn and Tickle, 1999). Furthermore, the AER of the dolphin and of the lizard *Anguis fragilis* are transient structures, which indicates that the failure in maintenance of AER-derived signaling could account for the degeneration of the limb bud. In contrast, the python does not have a morphologically distinct AER nor expresses the genes normally associated with this signaling center (FGF, Dlx, Msx; Cohn and Tickle, 1999), but the limb bud does not degenerate. Shh is also absent from the forelimb buds of *Calyptommatus*, but further investigation is needed in order to determine the causative factors behind limb bud degeneration.

## Conclusions

Considering the great diversity of limbs, it seems equally likely that there exists a corresponding diversity of developmental mechanisms. We are still distant from a comprehensive understanding of the anatomy and evolutionary relationships among species, and we are much farther from understanding the mechanisms by which different limbs are formed, especially considering the dynamic properties of developmental systems. Minor limb reductions can be the result of diverse changes in the developmental program, such as differential regulation of specific signals or of its receptors. However, in greatly reduced limbs, the major signaling pathways (FGF and Shh signaling) seem to be consistently affected, although these analyses do not go further into the dissection of the pathways and its regulations due to the experimental challenges of working with non-model organisms. This apparent convergence regarding the involvement of these major patterning signals as the potential drivers of limb loss can be nothing more than the result of the relaxation of the selective pressures over these key signaling pathways after long evolutionary times.

## Acknowledgements

Our work was supported by Conselho Nacional de Pesquisa (CNPq) and Fundação de Amparo à Pesquisa do Estado de São Paulo (FAPESP). JGR is truly grateful to Cheryll Tickle for helping in establishing the whole mount in situ. We are also greateful for the reviewer’s constructive comments.

## Frequent abbreviations

d=: digit
mc=: metacarpal
Shh=: sonic hedgehog
AER=: apical ectodermal ridge
Fgf=: fibroblast growth factor
Bmp=: bone morphogenetic protein
Wnt=: wingless homologue gene

## Supplementary information

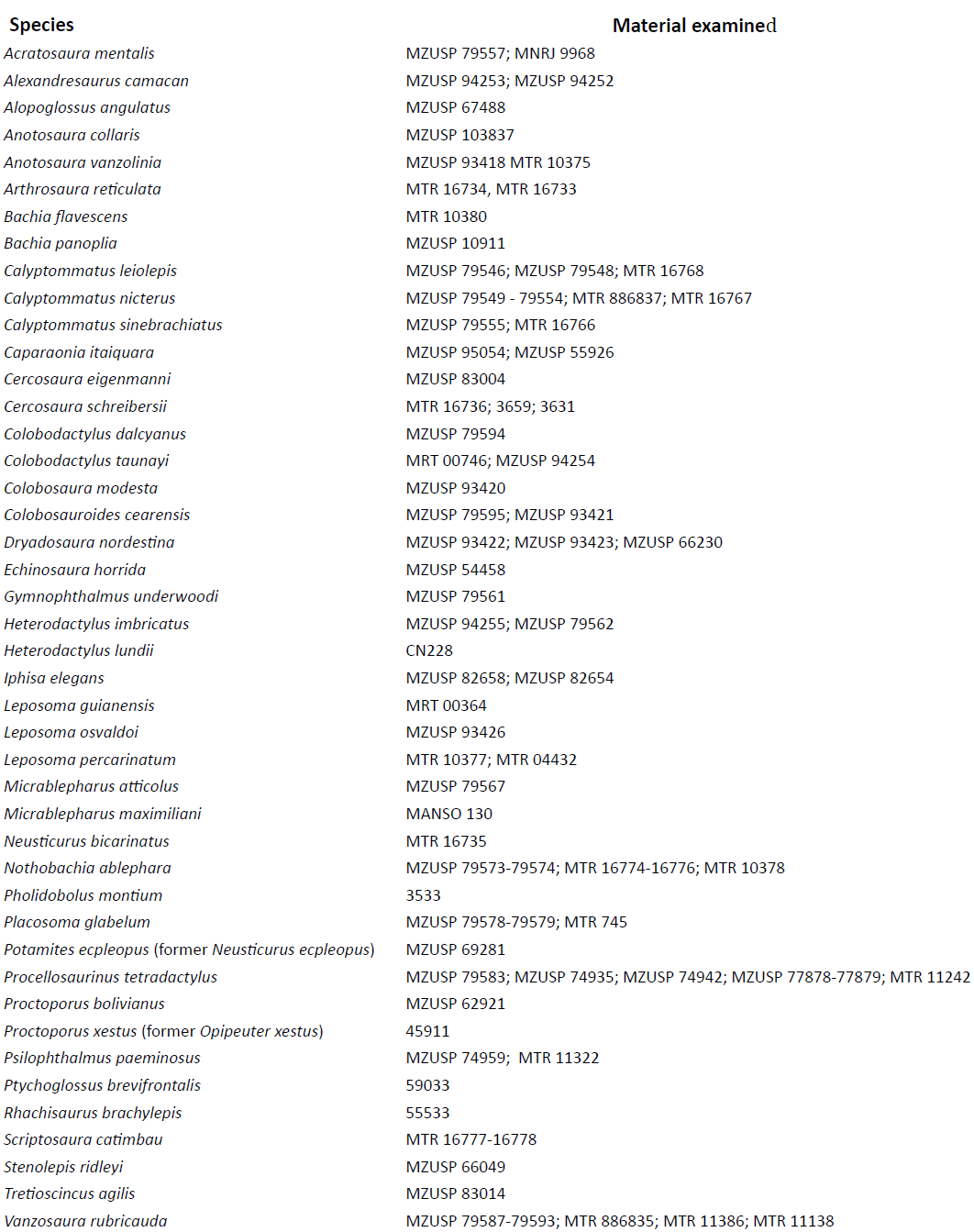

